# WT1 targeting multi-epitope vaccine design for glioblastoma multiforme using immuno-informatics approaches

**DOI:** 10.1101/2024.06.19.599735

**Authors:** Tanvir Ahmed

## Abstract

Glioblastoma multiforme (GBM) is a highly aggressive form of brain cancer classified as grade 4 glioma with a median survival rate of 12-14 months. Currently there is no cure and the conventional treatment outcomes are poor making it imperative to develop novel treatments. WT1, also known as Wilms’ Tumor 1, is a protein that is often found to be overexpressed in GBM and has minimal expression in normal tissues has become a promising target for immunotherapy for its oncogenicity and immunogenicity. This study aimed to develop a multi-epitope vaccine using immuno-informatics approaches that specifically targets the WT1 protein. The WT1 sequence was used to predict B and T cell epitopes which showed probable antigenic, non-allergic and non-toxic properties. Using suitable linker and adjuvants the vaccine construct (370 amino acids) was prepared which were then analyzed for solubility, physicochemical properties, molecular docking to show receptor interactions and molecular dynamics to show stability of the vaccine-receptor complexes. Subsequently, the vaccine sequence was back translated (1110 nucleotides), codon adaptation, pET-28a (+) vector in-silico cloning, and immune response simulations were performed. The designed vaccine lays groundwork for future in-vitro and in-vivo studies and potentially develop it into a novel treatment option for GBM patients.

## 1. Introduction

Glioblastoma multiforme (GBM) is the most common type of primary malignant brain tumor in adults. It develops from glial or precursor cells. [1, 2]. Even with extensive medical interventions such as surgery, chemotherapy, and radiation therapy, the average life expectancy remains limited to a range of 12–14 months. [3–5]. Extensive research has been conducted on the molecular mechanisms of glioma cell tumorigenicity, but significant clinical progress has yet to be made. [6, 7]. Researchers are currently studying ways to improve therapeutic approaches by gaining a more comprehensive understanding of the pathophysiology associated with GBM development. [1]. Continued research is currently underway to explore novel therapies for recurring GBM. [8–10], with immunotherapies showing promise [11].

Immunotherapy harnesses the inherent ability of the immune system to combat various diseases. Unlike conventional therapies, immunotherapy works by boosting the body’s immune system’s ability to recognize and eliminate cancerous cells [12, 13]. This approach offers several advantages, including the potential for long-term immunological memory, reduced side effects that are systemic, and the ability to tackle the challenge of tumor heterogeneity. [12]. Immunotherapy mediated by antibodies and based on dendritic cells are two therapeutic approaches used to combat malignant gliomas [14]. Another form of treatment that shows promise is cancer vaccination. [15]. Therapeutic cancer vaccines have demonstrated significant potential in eliciting preventive and therapeutic immune responses against GBM, among other immunotherapy approaches. Vaccines expose the immune system to antigens, which help it identify and eliminate cancer cells that present those molecules. This proactive approach not only focuses on existing cancers but also has the potential to prevent the development or return of tumors. Recent progress in the fields of molecular biology and tumor immunology has resulted in the identification of multiple tumor-associated antigens. These antigens have shown promise for cancer vaccination, as their epitopes, which are connected to HLA Class I molecules, have been recognized by CTLs. Selecting the appropriate target antigen is of utmost importance when developing an effective cancer vaccine. To achieve optimal performance, the target antigen must possess two essential characteristics: This phenomenon is observed in tumors with high expression, where the immune response is targeted towards cancer cells while minimizing damage to healthy tissues. The antigen must be highly immunogenic to elicit a robust immune response that includes both humoral and cellular responses. WT1, the product of the Wilms tumor gene, has been recognized as one of the tumor-associated antigens. [15, 16].

The identification of Wilms’ tumor 1 (WT1) as a possible oncogene in glioblastomas and other cancers, including those arising from the breast and hematological system, has been established. [3, 17–20]. WT1 expression is observed in around 80% of glioblastoma specimens and over 50% of malignant astrocytoma cell lines [21]. WT1 has demonstrated promise as a therapeutic target due to its capacity to regulate various facets of tumorigenesis. Furthermore, the healthy brain does not exhibit any signs of its presence [3, 21–23]. The WT1 gene has been identified as the gene linked to Wilms tumor. The gene codes for a zinc finger transcription factor involved in cell division, proliferation, apoptosis, and organ development, among other biological processes. The WT1 gene was initially classified as a tumor-suppressor gene, but subsequent research indicated that it may function as an oncogene rather than a tumor-suppressor gene [15]. The WT1 gene, located at chromosome locus 11p13, has been found to encode a polypeptide ranging in length from 52 to 55 kilobases. This polypeptide possesses four zinc fingers at its C-terminal & an N-terminal transactivation region [21, 24]. WT1 is widely recognized as a highly promising target for cancer immunotherapy, owing to its specificity, oncogenicity, immunogenicity, as well as therapeutic significance [25, 26]. The WT1 protein triggers both humoral [27, 28] and cellular immunological responses in-vitro [29–31] and within in-vivo [32]. Extensive research has resulted in the advancement of cancer immunotherapies that specifically focus on WT1, such as vaccines based on peptides [33–35], WT1 mRNA-electroporated dendritic cell (DC) therapy [36], and WT1 peptide-pulsed DC therapy, which have shown clinical effectiveness in different types of malignant tumors [37, 38].

WT1 shows potential as a target, however, the intertumoral heterogeneity of GBM poses a significant challenge. The presence of different WT1 epitopes within tumor cells in the same GBM has been observed. These epitopes are short peptide fragments that are recognized by the immune system. A vaccine that focuses on a single epitope may only work against a particular set of tumor cells. This could result in the development of resistant clones, which can contribute to the reoccurrence of the disease.

Multi-epitope vaccines have emerged as a powerful solution to address this challenge. The vaccines include epitopes from various sections of the target antigen, encompassing a broad spectrum of MHC binding specificities. The increased range of epitopes in the vaccine allows it to stimulate an immune response against a wider array of GBM cell types, potentially addressing tumor heterogeneity and enhancing treatment efficacy.

The objective of this study is to develop a multi-epitope vaccine for GBM by employing immuno-informatics techniques, with a specific emphasis on the WT1 protein. The in-silico vaccine model aimed to predict B and T cell binding epitopes predicted antigenicity and simulating immune responses against GBM. Its lays a promising foundation for future potential experimental studies to develop novel GBM immunotherapy.

## 2. Methods

### 2.1 Protein sequences identification, retrieval, and expression analysis

The protein sequence of WT1, specifically the isoform E of Wilms tumor protein in Homo sapiens, was obtained from the National Centre for Biotechnology Information (NCBI). (https://www.ncbi.nlm.nih.gov/). The selected sequence is in the FASTA format and corresponds to Wilms tumor protein isoform E in Homo sapiens. It has a length of 302 amino acids and is identified by the NCBI Reference Sequence NP_001185480.1. The graphics option was selected to visualize the structural components of the retrieved sequence. Then, for expression analysis, the UALCAN (https://ualcan.path.uab.edu/index.html)[39, 40] database TCGA option was used to determine the expression of WT1 in GBM based sample types and KM plot was derived from the survival analysis of WT1 based on race.

### 2.2 Prediction of B and T cell epitopes

The prediction of T-cell and B-cell epitopes was conducted using the Immune Epitope Database (IEDB) (https://www.iedb.org/), which contains a vast amount of experimental data on antibodies and epitopes. The epitope analysis resource was used to select the B cell epitope prediction option. The FASTA protein sequence was then pasted for the prediction of linear epitopes from the protein sequence. The Bepipred Linear Epitope Prediction 2.0 method was chosen and subsequently submitted. Afterwards, the Peptide binding to MHC class I molecules and Peptide binding to MHC class II molecules tool were used to predict the T cell epitopes for MHC binding. The IEDB NetMHCPan 4.1 EL (epitope prediction) prediction method was utilized for MHC I. The peptides were sorted based on their predicted score, and only the epitopes with a score of 0.9 were chosen for further analysis. Afterwards, the NetMHCIIPan 4.1 EL (epitope prediction) was utilized for MHC II as recommended. The peptides were sorted based on their adjusted rank, and only those with a rank less than 1 were chosen for further analysis.

### 2.3 Antigenicity and allergenicity of the predicted epitopes

Before proceeding with vaccine design, the antigenicity and allergenicity of each predicted epitope (both B and T cell) were thoroughly tested. The antigenicity assessment tool VaxiJen v2.0 was utilized to evaluate the antigenicity of the epitopes. The tool can be accessed at https://www.ddg-pharmfac.net/vaxijen/VaxiJen/VaxiJen.html. A tumor model was chosen with a threshold of 0.5. If an epitope has an antigenicity score above 0.5, it is deemed worthy of consideration as a potential antigen and undergoes further evaluation. In addition, the epitopes underwent allergenicity testing using the AllerTOP v. 2.0 (https://www.ddg-pharmfac.net/AllerTOP/). Epitopes that are chosen for further evaluation are those that are likely to be non-allergenic.

### 2.4 Prediction of the epitopes’ ability to induce cytokines

Determining the induction capacity of the predicted HTL epitopes for interferon-γ (IFN-γ) was done using the IFNepitope (http://crdd.osdd.net/raghava/ifnepitope/). The Motif and SVM hybrid approach was used and the IFN-gamma versus other cytokine model was employed for predicting the IFNepitopes. Similarly, the IL-10Pred (http://crdd.osdd.net/raghava/IL-10pred/) was used to predict the IL-10 inducing HTL epitopes. The SVM prediction model was used with the default SVM threshold set at -0.3.

### 2.5 Construction of multi-epitope vaccine sequence

Using an adjuvant and linkers, the most promising antigenic epitopes were joined to form a fusion peptide. Based on the predicted B-cell and T-cell epitopes, the candidate vaccine sequence was created by fusing the linear B- and T-cell epitopes with linkers. The linkers used in vaccine construction were: EAAAK, GPGPG, AAY. The adjuvant used was Cholera Toxin Subunit B (GenBank: AND74811.1) to enhance the vaccine’s ability to trigger the innate immune response [41]. The length of the peptide vaccine design was 370 amino acids. An analysis of the vaccine sequence’s antigenicity, allergenicity, and toxicity was conducted using VaxiJen v2.0 (https://www.ddg-pharmfac.net/vaxijen/VaxiJen/VaxiJen.html), AllerTOP v. 2.0 (https://www.ddg-pharmfac.net/AllerTOP/) [42], and Toxinpred (http://crdd.osdd.net/raghava/toxinpred/) [43], respectively.

### 2.6 Physicochemical property analysis

The online web server ProtParam (https://web.expasy.org/protparam/) [44] was used to determine several physicochemical characteristics of the candidate peptides, such as theoretical pI, aliphatic index, instability index, estimated half-life in mammalian reticulocytes in vitro, extinction coefficient, molecular weight, and grand average of hydropathicity (GRAVY). An estimation of the vaccine peptide’s solubility in water was conducted using the Pepcalc (https://pepcalc.com/) tool [45]. The physiochemical properties predicted include number of residues, molecular weight, extinction coefficient, iso-electric point, net charge at pH 7, and estimated solubility.

### 2.7 Prediction of the secondary and tertiary structure of the vaccine construct

The vaccine construct’s secondary structure was predicted using the PSIPRED tool (PSIPRED 4.0) (http://bioinf.cs.ucl.ac.uk/psipred/) [46], the SIMPA96 tool (https://npsa-prabi.ibcp.fr/cgi-bin/npsa_automat.pl?page=/NPSA/npsa_simpa96.html) [47], SOPMA (https://npsa-pbil.ibcp.fr/cgi-bin/npsa_automat.pl?page=/NPSA/npsa_sopma.html) [47] with the parameters: 4 conformational states, 8 similarity thresholds, and 17 window widths. These servers are reliable, rapid, and effective for predicting the percentages or quantities of amino acids in α helix, β-sheet, and coil structural formations [48–51]. The tertiary structure was predicted using the SCRATCH Protein Predictor’s 3Dpro tool (https://scratch.proteomics.ics.uci.edu/).

### 2.8 Tertiary structure refinement and validation of the vaccine

The predicted vaccine 3D model was improved in resolution to more closely reflect the native protein structure by employing the GalaxyWEB server’s GalaxyRefine module (https://galaxy.seoklab.org/cgi-bin/submit.cgi?type=REFINE) [52–54]. To refine the tertiary protein structures, the server uses dynamic simulation and CASP10 tests the refinement strategy. In addition, the refined protein was validated by analyzing the Ramachandran plot generated by PROCHECK (https://saves.mbi.ucla.edu/) [55, 56]. In addition to PROCHECK, another web platform called ProSA-web (https://prosa.services.came.sbg.ac.at/prosa.php) [57, 58] was also used for protein validation. ProSA-web validates protein tertiary structures and calculates a 3D-structure quality score. If the score falls outside the range of native proteins, the structure may contain errors. A z-score indicating the overall model quality of a query protein structure is produced by the ProSA-web.

### 2.9 Molecular docking of the vaccine candidate

The vaccine protein was docked against the MHC class I (PDB ID: 1I1Y) receptor, and the MHC class II (PDB ID: 1KG0) and against the TLR4 (PDB ID: 4G8A) receptor using the CLUSPRO 2.0 (https://cluspro.bu.edu/login.php) [59–62] protein-protein docking server. The software uses three different methods. The first method involves a fast Fourier transform (FFT) correlation approach. The second method involves grouping the best energy conformations based on root-mean-square deviation (RMSD). Lastly, the software evaluates the stability of clusters. The ClusPro 2.0 server uses PIPER for sampling. The center of mass of the receptor is situated at the origin of the coordinate system, while the positions of the ligand’s rotation and translation are determined using a specified level of discretization. The rotational space is sampled using a grid that is based on a sphere. The spherical surface is subdivided in such a way that each pixel covers an equal surface area. Around 70,000 rotations correspond to approximately 5 degrees in Euler angles. The translational grid has a step size of 1 Å. To adequately explore the conformational space of an average-sized protein, a substantial number of conformations, ranging from 109 to 1010, must be sampled. The expression E = w1E_rep_ + w2E_attr_ + w3E_elec_ + w4E_DARS_ represents the interaction energy between two proteins. In this context, E_rep_ and E_attr_ denote the repulsive and attractive components of the van der Waals interaction energy, respectively, while E_elec_ refers to the electrostatic energy term. The E_DARS_ method refers to a structure-based potential called the DARS29 approach, which is commonly known as Decoys as the Reference State. The study primarily examines the impact of removing water molecules from the interface, specifically the resulting free energy change due to desolvation. The coefficients w1, w2, w3, and w4 are selected to represent the weights of specific terms in different docking scenarios. [61].

### 2.10 Molecular dynamics simulation of the docked complexes

The i-MOD server (i-MODS) (https://imods.iqfr.csic.es/) was used for conducting molecular dynamics simulation for the complexes of the vaccine candidate with the MHC I, MHC II, TL4 receptors. An evaluation of the protein binding stability as well as minimal deformation of the docked complex was conducted using normal mode analysis (NMA) with i-MODS. The NMA uncovers the inherent movement of the complex coordinates. The protein’s stability is illustrated by its main-chain deformability plot, which assesses the binding stability and potency of the ligand to the receptor using factors like atomic fluctuations, B-factor values, covariance matrix, elastic network model, & eigenscore. The eigenscore provides insight into the complex’s motion rigidity and can be compared with deformation profiles. The atomic coordinates can be assessed for stability and susceptibility to deformation by utilizing independent component algorithms. A low Eigen score is indicative of reduced stability [41, 63].

### 2.11 Back translation of the vaccine and codon adaptation

Back translation involves converting an amino acid sequence into a DNA sequence. The amino acid sequence of the vaccine construct is converted to nucleic acid sequence using EMBOSS Backtranseq (https://www.ebi.ac.uk/jdispatcher/st/emboss_backtranseq) [64]. EMBOSS Backtranseq is a tool used to predict the nucleic acid sequence corresponding to a given protein sequence. The codon usage table is used to determine the frequency of each codon’s usage for every amino acid. To express a foreign gene in a host organism, it is required to optimize the codons based on the specific host organism. The codons in the vaccine construct were optimized based on E. coli K12 in the JCat server (https://www.jcat.de/) [65] to enhance translation efficiency. Optimizing codons increases the expression rate of the final vaccine in E. coli K12 as an expression host due to differences between human and host codons [41]. The peptide vaccine construct consisting of 370 amino acid residues generated 1110 nucleotide sequences. An analysis was conducted on the optimized codon sequence to determine its expression parameters, including the codon adaptation index (CAI) and the percentage of GC-content. The ideal CAI value is 1.0, with a score above 0.8 being regarded acceptable. The optimal GC content is between the range of 30 to 70% [66, 67].

### 2.12 In silico cloning of the vaccine

Using the SnapGene (https://www.snapgene.com/free-trial) restriction cloning module, the DNA sequence of the codon-adapted vaccine was introduced into the pET-28a(+) vector plasmid between the BlpI and BSSHII restriction sites to ensure vaccine expression.

### 2.13 Immune response simulation

Immune simulations were conducted using the C-ImmSim server (http://kraken.iac.rm.cnr.it/C-IMMSIM/) to assess the immune response profile of the constructed multi-epitope vaccine. The C-ImmSim is an agent-based computational tool that uses a position-specific score matrix (PSSM) and machine learning techniques to predict epitope and immune interactions [68, 69]. The simulation was conducted using the default parameters. The simulation involved a volume of 1,000 units, 1,000 simulation steps, a random seed of 12,345, and the administration of a vaccine without LPS [41, 70, 71].

## 3. Results

### 3.1 WT1 sequence retrieval and expression analysis

The Wilms tumor protein isoform E [Homo sapiens] (**Figure 1**) having an accession number of NP_001185480.1 is of 302 amino acids length. The FASTA sequence of this protein is used for the vaccine design. The sequence is as follows: MEKGYSTVTFDGTPSYGHTPSHHAAQFPNHSFKHEDPMGQQGSLGEQQYSVPPPVYGCHTPTDSCTGSQALLLRTPYSSDNLYQMTSQLECMTWNQMNLGATLKGVAAGSSSSVKWTEGQSNHSTGYESDNHTTPILCGAQYRMHTHGVFRGIQDVRRVPGVAPTLVRSASETSEKRPFMCAYPGCNKRYFKLSHLQMHSRKHTGEKPYQCDFKDCERRFSRSDQLKRHQRRHTGVKPFQCKTCQRKFSRSDHLKTHTRTHTGEKPFSCRWPSCQKKFARSDELVRHHNMHQRNMTKLQLAL. **(Figure 1A)** Wilms tumor protein isoform E features, region features, and site features. The protein characteristics of isoform E of the Wilms tumor protein are displayed in this section of the figure. It encompasses areas with characteristics such as C2H2 Zn finger and COG5048. One kind of protein domain that aids in a protein’s ability to bind to DNA is the C2H2 Zn finger. This shows that Wilms tumor protein isoform E might be involved in the control of gene expression. The protein may bind to nucleic acids because it has a putative nucleic acid binding site. **(Figure 1B)** WT1 Expression in GBM according to Sample Types. This section demonstrates that compared to samples of normal brain tissue, initial tumor samples have higher levels of WT1 expression. **(Figure 1C)** The impact of race and WT1 expression level on the survival of GBM patients. This section of the figure examines the KM plot of race, WT1 expression level, and GBM patient survival rate. Based on race and WT1 expression level, there is no statistically significant variation in the survival rate.

**Figure 1:**
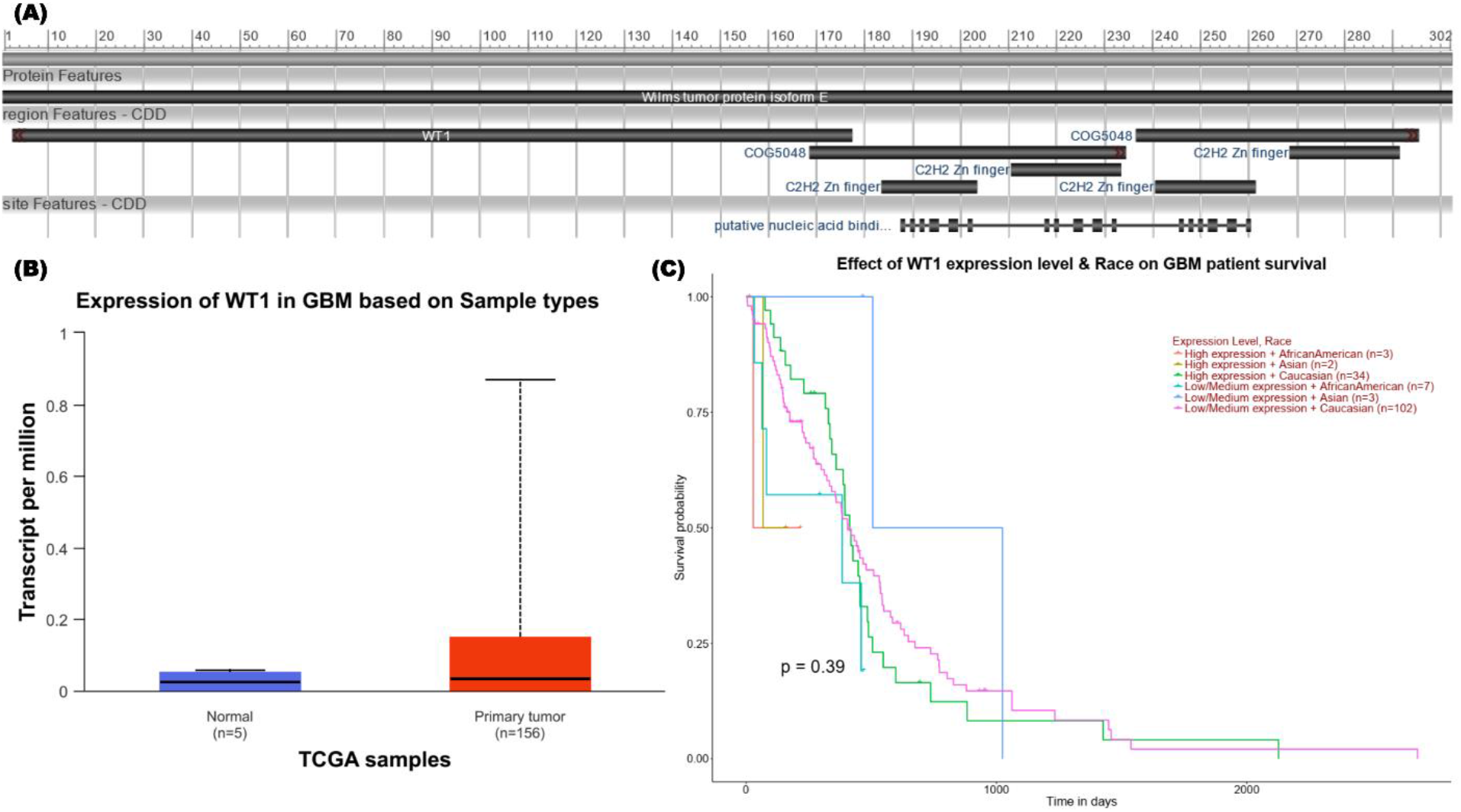
The Wilms tumor protein **(A)** isoform E features, region features and site features. **(B)** Expression of WT1 in GBM based on sample types **(C)** Effect of WT1 expression level and race on GBM patient survival.

### 3.2 Selecting the most promising B and T cell epitopes

At initial analysis, the protein is found to have probable antigenic properties (0.5418). Following this, the WT1 protein was chosen as a model for predicting T-cell (MHC class I and II) and B-cell epitopes using the IEDB server to develop the multi-epitope vaccine. Based on the ranking, the top-scoring MHC Class-I or Cytotoxic T Lymphocyte (CTL) and MHC Class-II or Helper T Lymphocyte (HTL) epitopes, as well as one B-cell epitope, were chosen for further evaluation. Following this, a select few criteria were used to evaluate the top epitopes, including strong antigenicity, lack of allergenicity, and non-toxicity. Furthermore, the researchers evaluated the ability of HTL epitopes to stimulate cytokines like IFN-γ, IL-4, and IL-10 to assess their immunogenicity. In the end, epitopes that met these criteria were singled out as the most promising as listed in **Table 1**. The B cell epitope selected was SSSVKWTEGQSNHSTGYESDNHTTPILCGAQYRMHTHGVFRGIQDVRRVPGV. For finalizing the T cell epitopes, MHC I binding epitopes were: SASETSEKR, TSQLECMTW, GQSNHSTGY, MTSQLECMTW, MTSQLECMTW, VTFDGTPSY and MHC II binding epitopes were: APTLVRSASETSEKR, PGVAPTLVRSASETS, RVPGVAPTLVRSASE, VPGVAPTLVRSASET, LRTPYSSDNLYQMTS, RTPYSSDNLYQMTSQ.

**Table 1:** Predicted B and T cell epitopes finalized for the candidate vaccine construction.

| T cell MHC I | Epitope | allele sequence | Length | Score Percentile | Rank | Antigenicity Score | Antigenicity | Allergenicity | Toxicity |
| --- | --- | --- | --- | --- | --- | --- | --- | --- | --- |
|  | SASETSEKR | HLA-A*68:01 | 9 | 0.907999 | 0.07 | 1.1416 | Antigen | Non-Allergen | Non-Toxin |
|  | TSQLECMTW | HLA-B*57:01 | 9 | 0.932306 | 0.08 | 1.0966 | Antigen | Non-Allergen | Non-Toxin |
|  | GQSNHSTGY | HLA-B*15:01 | 9 | 0.935303 | 0.01 | 1.0417 | Antigen | Non-Allergen | Non-Toxin |
|  | MTSQLECMTW | HLA-B*57:01 | 10 | 0.983967 | 0.02 | 1.0122 | Antigen | Non-Allergen | Non-Toxin |
|  | MTSQLECMTW | HLA-B*58:01 | 10 | 0.962671 | 0.03 | 1.0122 | Antigen | Non-Allergen | Non-Toxin |
|  | VTFDGTPSY | HLA-B*35:01 | 9 | 0.948628 | 0.02 | 0.5801 | Antigen | Non-Allergen | Non-Toxin |
|  | VTFDGTPSY | HLA-A*30:02 | 9 | 0.930595 | 0.01 | 0.5801 | Antigen | Non-Allergen | Non-Toxin |
|  | VTFDGTPSY | HLA-B*15:01 | 9 | 0.923911 | 0.01 | 0.5801 | Antigen | Non-Allergen | Non-Toxin |
|  | VTFDGTPSY | HLA-A*26:01 | 9 | 0.908753 | 0.02 | 0.5801 | Antigen | Non-Allergen | Non-Toxin |
| T cell MHC II | APTLVRSASETSEKR | HLA-DRB1*04:01 | 15 | 0.7582 | 0.96 | 1.0625 | Antigen | Non-Allergen | Non-Toxin |
|  | PGVAPTLVRSASETS | HLA-DQA1*01:02/DQB1*06:02 | 15 | 0.7209 | 0.29 | 0.7308 | Antigen | Non-Allergen | Non-Toxin |
|  | RVPGVAPTLVRSASE | HLA-DQA1*01:02/DQB1*06:02 | 15 | 0.81 | 0.08 | 0.6564 | Antigen | Non-Allergen | Non-Toxin |
|  | VPGVAPTLVR<br>SASET | HLA-DQA1*01:02/D<br>QB1*06:02 | 15 | 0.8354 | 0.07 | 0.6287 | Antigen | Non-Allergen | Non-Toxin |
|  | L RTPYSSDNL<br>YQMTS | HLA-DRB3*01:01 | 15 | 0.5067 | 0.75 | 0.5822 | Antigen | Non-Allergen | Non-Toxin |
|  | RTPYSSDNL<br>YQMTSQ | HLA-DRB3*01:01 | 15 | 0.5618 | 0.77 | 0.5098 | Antigen | Non-Allergen | Non-Toxin |
|  | RTPYSSDNL<br>YQMTSQ | HLA-DQA1*05:01/D<br>QB1*02:01 | 15 | 0.5228 | 0.94 | 0.5098 | Antigen | Non-Allergen | Non-Toxin |
| **B Cell** | SSSVKWTEGQ<br>SNHSTGYESD<br>NHTTPILCGAQ<br>YRMHTHGVFR<br>GIQDVRRVPG<br>V | - | 52 | - | - | 0.76 | Antigen | Non-Allergen | Non-Toxin |

### 3.3 Vaccine construction and analysis of physicochemical properties

By attaching suitable linkers to the most promising T-cell and B-cell epitopes, the vaccine sequence was constructed. Furthermore, the HHHHHH sequence and Cholera Toxin Subunit B adjuvant were conjugated at the proper positions. The vaccine sequence is: MIKLKFGVFFTVLLSSAYANGTPQNITDLCAEYHNTQIHTLNDKIFSYTESLAGKREMAIITFKNGATFQVEVPGSQHIDSQKKAIERMKDTLRIAYLTEAKVEKLCVWNNKTPHAIAAISMANEAAAKSSSVKWTEGQSNHSTGYESDNHTTPILCGAQYRMHTHGVFRGIQDVRRVPGVGPGPGSASETSEKRAAYTSQLECMTWAAYGQSNHSTGYAAYMTSQLECMTWAAYVTFDGTPSYGPGPGAPTLVRSASETSEKRGPGPGPGVAPTLVRSASETSGPGPGRVPGVAPTLVRSASEGPGPGVPGVAPTLVRSASETGPGPGLRTPYSSDNLYQMTSGPGPGRTPYSSDNLYQMTSQHHHHHH. The vaccine sequence has a length of 370 amino acids. **Figure 2** illustrates the sequence and construct. A further analysis of the vaccine’s physicochemical properties revealed that it demonstrates possible antigenic characteristics with a likelihood of 0.5781. Additionally, it has been predicted to be non-allergic and non-toxic. The vaccine has a molecular weight of 39488.11 Daltons, a Theoretical isoelectric point (pI) of 8.48, a molecular formula of C_1725_H_2676_N_494_O_542_S_15_, and is composed of a total of 5452 atoms. The extinction coefficients at 280 nm (46090 and 45840 M^-1^ cm^-1^) can be utilized to approximate the concentration of the protein solution at 0.1% (1 g/L), given a certain protein structure. The marginal disparity between the two values indicates the potential existence of disulfide linkages (cystines) that could have a minor impact on the protein’s structure. The estimated half-life data suggests that the protein exhibits moderate stability in mammalian cells, with a half-life of approximately 30 hours. In contrast, the protein shows potentially strong stability in yeast, with a half-life of more than 20 hours, and in E. coli, with a half-life of over 10 hours. Nevertheless, the in vitro conditions may not accurately replicate the in vivo milieu in which the vaccine operates. The protein’s instability index (II) of 43.63 indicates a general tendency towards instability. This is a possible issue regarding the storage and duration of effectiveness of vaccines. The protein exhibits a mildly hydrophobic nature, as seen by its aliphatic index of 59.38 and negative GRAVY score of -0.479. This could potentially impact the protein’s interactions with other molecules and alter its function within the vaccine. Although the protein may exhibit stability in certain settings such as yeast, the in vitro half-life and instability index give rise to worries over its overall stability. This could pose a significant disadvantage for the vaccination, particularly in terms of storage and shelf life.

**Figure 2:**
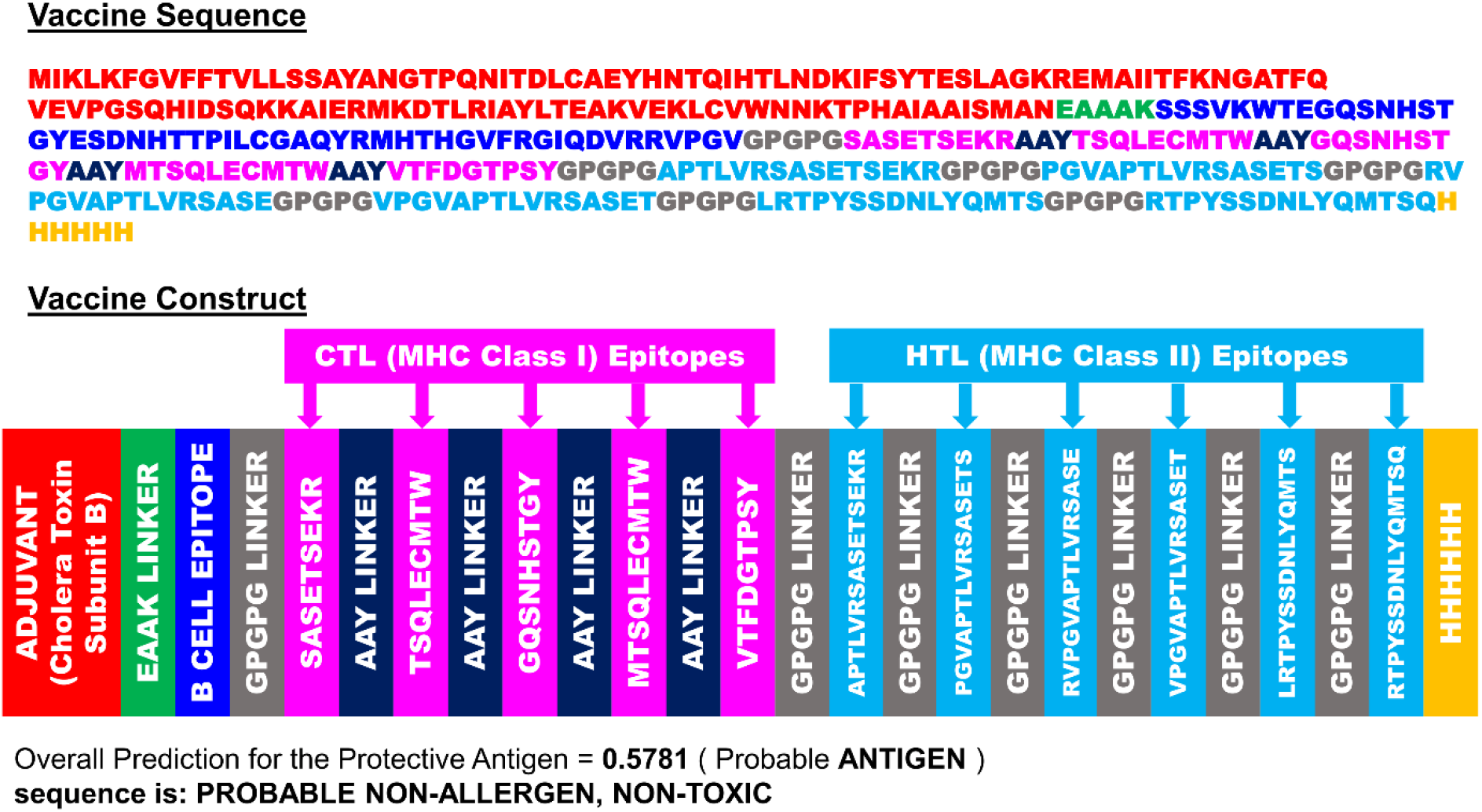
The designed vaccine amino acid sequence and the vaccine structure construct.

### 3.4 Secondary and tertiary vaccine structure prediction, refinement and validation

The data from SOPMA and SIMPA96 provides predictions on the secondary structure of the vaccine, offering valuable insights into its potential shape and function. Here’s a breakdown of the results: Both methods indicate a substantial amount (approximately 46-60%) of the vaccine is likely in a random coil conformation (Cc). Random coils provide flexibility and typically act as connectors between more organised sections. The distribution of secondary structures includes the alpha helix as the second most common structure at approximately 24-30%, with extended strands following at around 15-16%. Alpha helices and extended strands contribute to structural stability and frequently participate in interactions between proteins or functional regions. Major Contrasts: According to SOPMA, the percentage of random coil is slightly lower at 46.22% compared to SIMPA96’s prediction of 60.11%. It indicates that SOPMA anticipates a higher presence of structured elements. According to SOPMA, the percentage of alpha helix is slightly higher at 30.54% compared to SIMPA96’s 24.53%. Indicating that SOPMA anticipates a more inflexible arrangement. It is suggested that the vaccine may be flexible due to the high content of random coil, potentially offering advantages for antigen presentation. Implications of alpha helices and extended strands point to possible functional sites for interaction with the immune system or other molecules. The resuts of SOPMA and SIMPA96 analysis are listed in **Table 2**. The predicted secondary structure obtained from PSIPRED, SOPMA and SIMPA96 are illustrated in **Figure 3**. The tertiary structure was predicted using the SCRATCH protein predictor’s 3Dpro tool. The 3D structure of the protein was refined using the Galaxy WEB (**Table 3**). According to the data, the initial model achieved a GDT-HA score of 1.0000, showing a complete match with the reference structure. Although, it does have the highest RMSD (0.000) which could be attributed to overfitting. Models 1-5 demonstrate a decline in GDT-HA score (ranging from 0.9115 to 0.8973) in comparison to the original model, suggesting a slight decrease in resemblance to the reference structure. Yet, these models demonstrate a notable reduction in RMSD (ranging from 0.499 to 0.541), indicating an improved overall fit. The refined models (1-5) demonstrate a substantial enhancement in MolProbity score compared to the initial model, with a range of 1.725-1.837 versus 3.602. Implying improved geometry and reduced clashes. The Clash score further confirms this trend, as all refined models show significantly lower clash scores than the initial model. Poor rotamers are significantly reduced in the refined models, ranging from 0.3 to 1.0 compared to 5.0 in the initial model. The residues favoured by Rama consistently remain high in all models (above 88%), indicating a strong backbone conformation. All models show negative Z-scores, suggesting a level of energetic strain. Based on the data, it appears that the refined models (1-5) demonstrate a balance between similarity to the reference structure (GDT-HA) and overall fit (RMSD). Nevertheless, all models demonstrate notable enhancements in MolProbity score, clash score, and the number of poor rotamers, indicating superior model quality. MODEL 5 seems to be the top-performing model based on the combined weighted score of GDT-HA and MolProbity. This model strikes a fine balance between proximity to the reference structure (GDT-HA) and the general quality (MolProbity). For validation of the tertiary structure of Model 5, Ramachandran plot **(Figure 4)** was generated usisng the Procheck tool. The validity of the vaccine candidate’s tertiary structure is suggested by the Ramachandran plot for model 5, which shows that 91.7% of the residues are in the most favoured areas. This suggests that the protein’s backbone structure is probably stable and free of significant strain. Only 0.7% of the residues are in prohibited areas, while 7.6% of the residues are in additional permitted locations. This implies that there aren’t many potentially strained conformations in the structure overall, which makes it acceptable.

**Figure 3:**
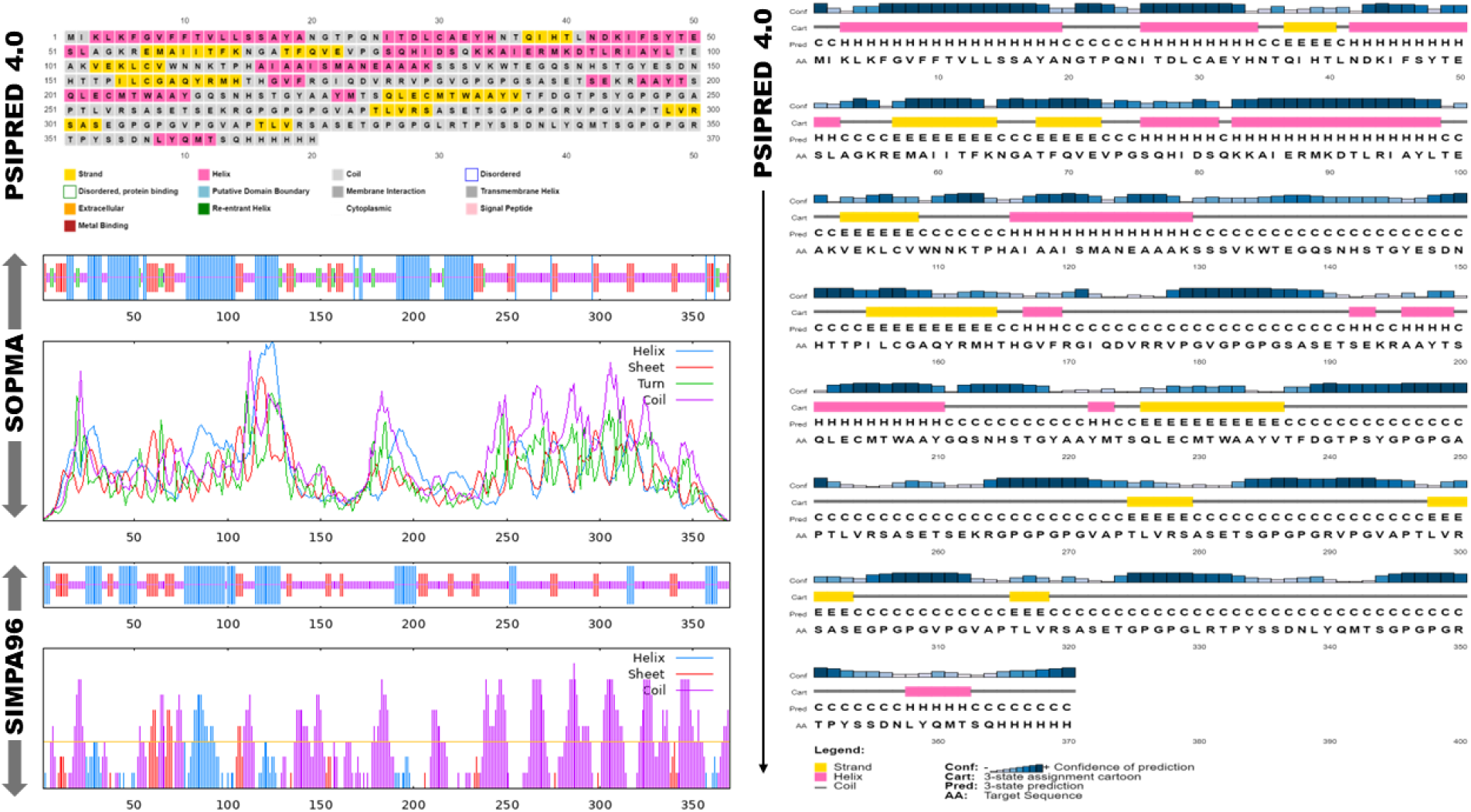
Secondary structure prediction of the vaccine using PSIPRED 4.0, SOPMA and SIMPA96

**Figure 4:**
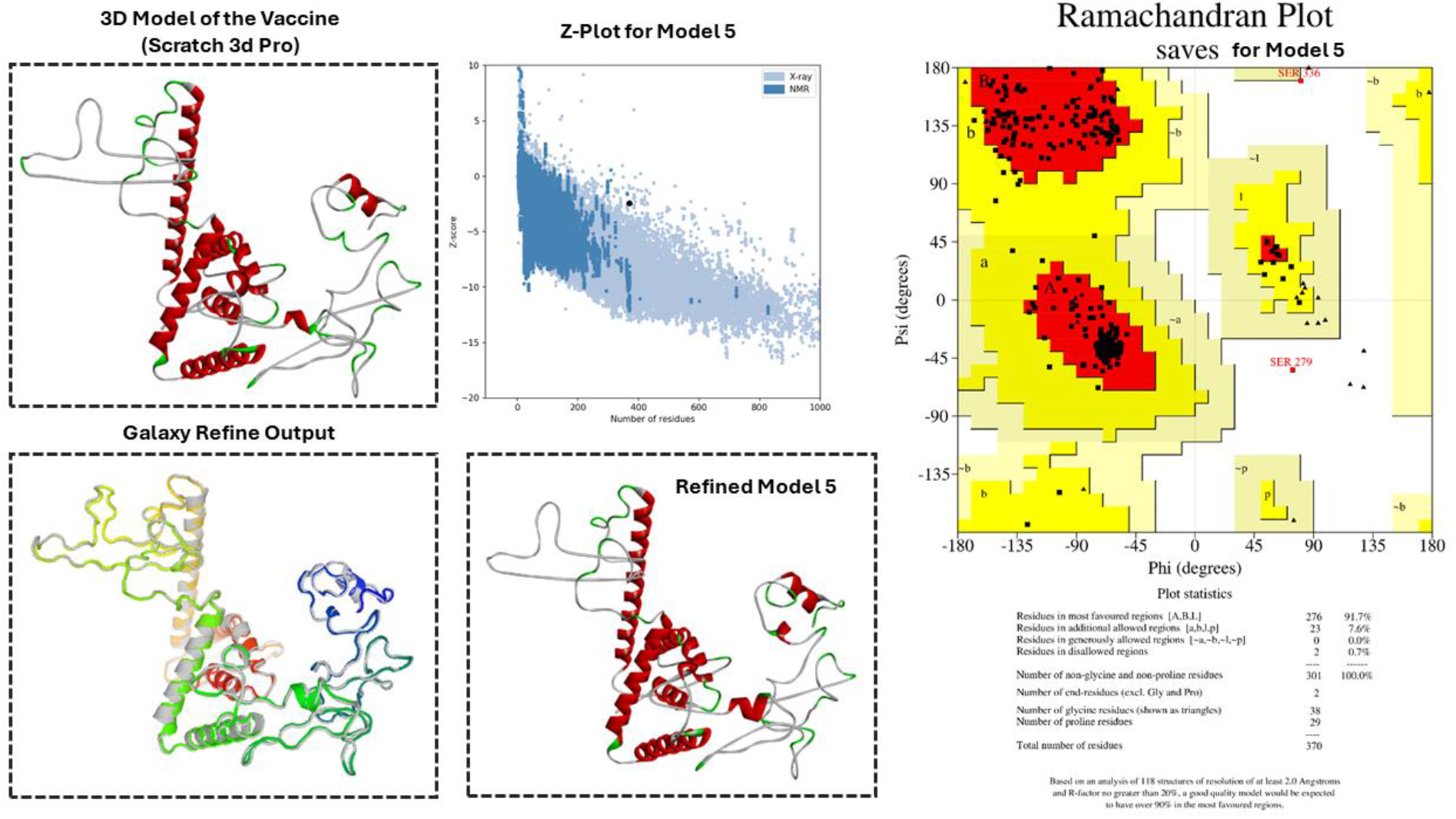
The 3D model (tertiary structure), refinement and validation (Z-Plot and Ramachandran Plot) of the vaccine structure

**Table 2:** The secondary strucure prediction of the vaccine from SOPMA and SIMPA96.

| Structure | Symbol | SOPMA |  | SIMPA96 |  |
| --- | --- | --- | --- | --- | --- |
|  |  | No. of Amino acids | Percentage | No. of Amino acids | Percentage |
| Alpha helix | Hh | 113 | 30.54% | 91 | 24.53% |
| 310 helix | Gg | 0 | 0.00% | 0 | 0.00% |
| Pi helix | Ii | 0 | 0.00% | 0 | 0.00% |
| Beta bridge | Bb | 0 | 0.00% | 0 | 0.00% |
| Extended strand | Ee | 61 | 16.49% | 56 | 15.09% |
| Beta turn | Tt | 25 | 6.76% | 0 | 0.00% |
| Bend region | Ss | 0 | 0.00% | 0 | 0.00% |
| Random coil | Cc | 171 | 46.22% | 223 | 60.11% |
| Ambiguous states | ? | 0 | 0.00% | 0 | 0.00% |
| Other states | - | 0 | 0.00% | 1 | 0.27% |

**Table 3:**
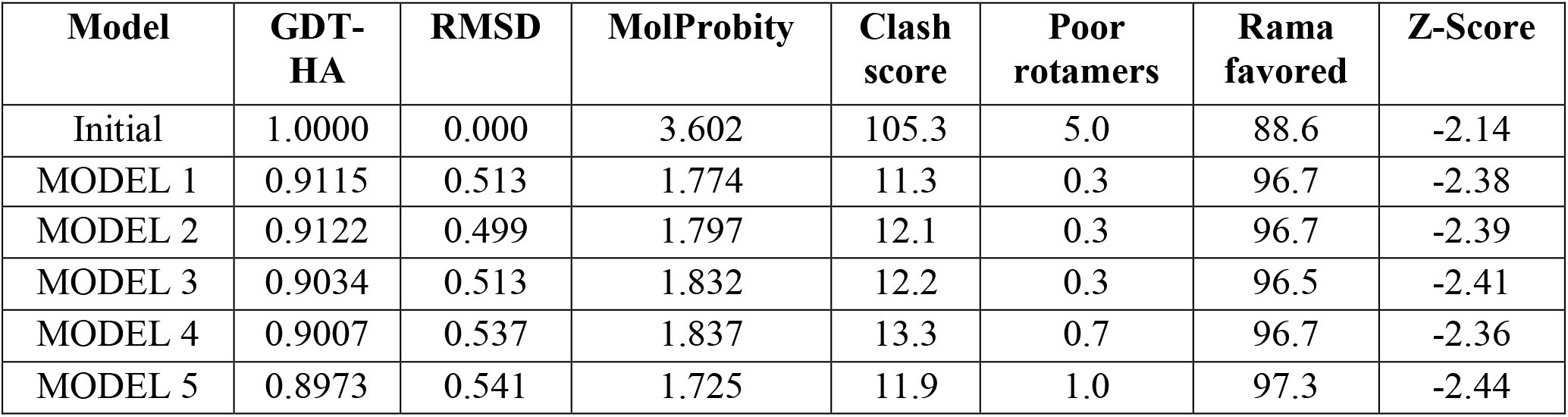
Tertiary structure refinement using Galaxy WEB.

### 3.5 Molecular docking of the vaccine

The results of protein-protein docking from CLUSPRO offered valuable insights into the interactions between receptors and vaccines. When comparing the three receptor types (MHC I, MHC II, and TL4), the analysis suggests that the MHC I receptor exhibits the strongest binding with the vaccine, with a weighted score of -1284.6. There appears to be a significant connection between the vaccine and MHC I receptor, suggesting a potent immune response triggered by the vaccine via MHC I presentation of antigenic peptides to cytotoxic T cells. While MHC II and TL4 receptors show binding with the vaccine, their weighted scores (-1205.4 and -1206.4, respectively) are higher than MHC I, suggesting weaker interactions (results of docking are presented in **Table 4**). The docked complexes are illustrated in **Figure 5**. Nevertheless, the variations in scores between MHC II and TL4 receptors are somewhat insignificant. Although the MHC II receptor has a lower weighted score, it exhibits lower energy in its center, suggesting a potentially more stable binding in that configuration.

**Figure 5:**
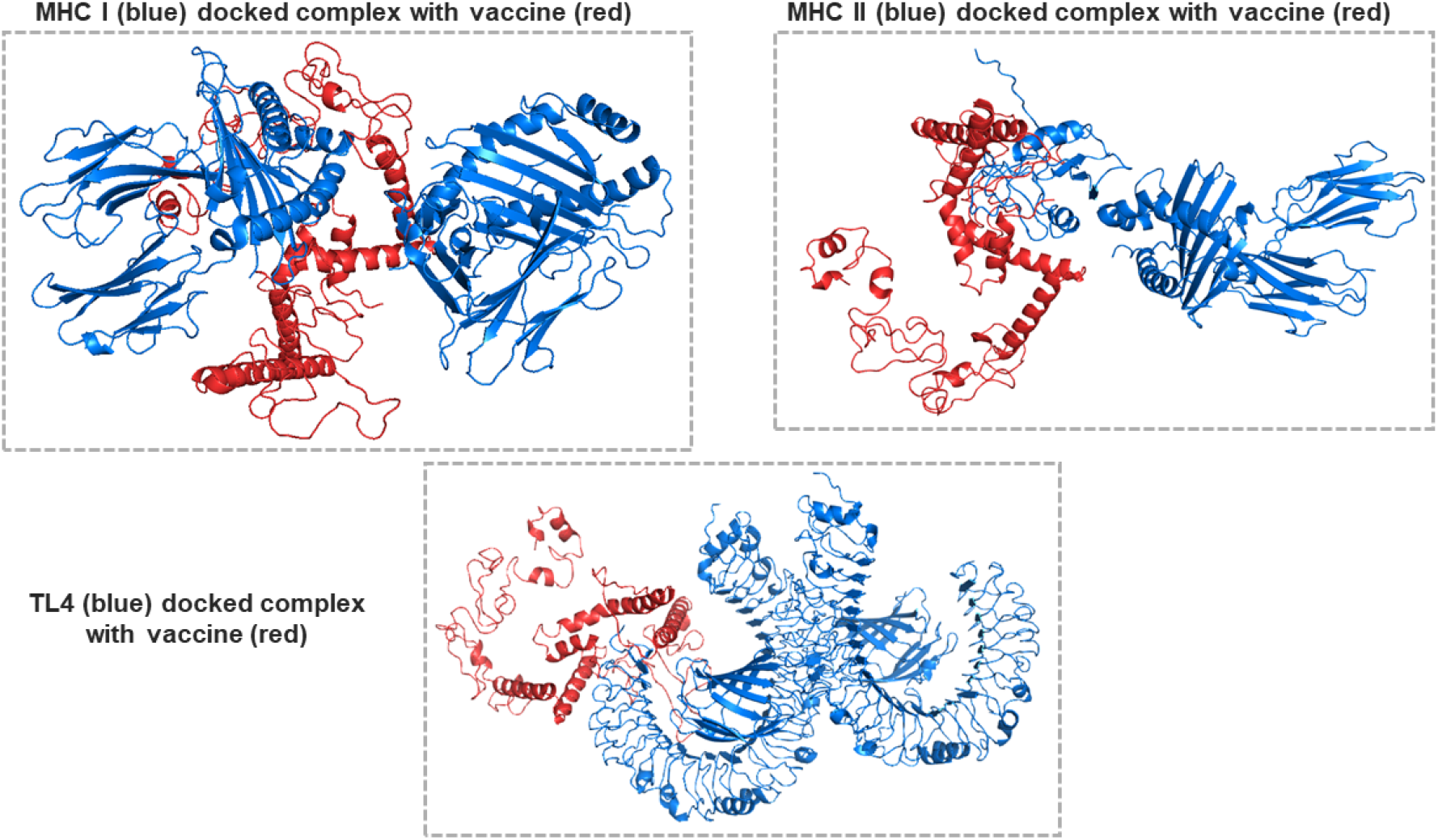
Molecular docked complexes of the vaccine and receptors (MHC I, MHC II and TL4)

**Table 4:** Vaccine-receptor complex docking results of model 0 from CLUSPRO.

| Receptor type in complex | Cluster | Members | Representative | Weighted Score |
| --- | --- | --- | --- | --- |
| MHC I | 0 | 42 | Center | -1174.6 |
|  |  |  | Lowest Energy | -1284.6 |
| MHC II | 0 | 49 | Center | -1108.9 |
|  |  |  | Lowest Energy | -1205.4 |
| TL4 | 0 | 50 | Center | -1051.2 |
|  |  |  | Lowest Energy | -1206.4 |

### 3.6 Molecular dynamics simulation of the docked complexes

An evaluation of the GBM candidate vaccine complex and receptor (MHC I, MHC II, and TL4) using the I-MODS molecular dynamics simulation revealed that MHC I exhibits the highest level of main-chain flexibility compared to all others. MHC II exhibits a similar pattern to MHC I, however with lower overall values. TL4 appears to demonstrate the least amount of flexibility in its main chain. The experimental B-factor for MHC I is higher than that of MHC II and TL4. The MHC I molecule demonstrates superior flexibility or mobility in comparison to the other two receptors. The eigenvalues of MHC I are the most elevated, subsequent to that of MHC II, and then TL4. Eigenvalues provide a quantitative measure of the overall motion exhibited by a molecule. Larger eigenvalues indicate greater flexibility. The variance plots of all three receptors display resemblances, with prominent peaks observed in the vicinity of residue 200 and 400. Variance measures the extent to which atoms deviate from their average positions. The covariance map and elastic network depict the interconnected movements of different components of the molecule. The results of the MD simulation are illustrated in **Figure 6**. Overall findings, The GBM vaccine complex engages with MHC I, MHC II, and TL4 in separate and specific ways. According to the data, MHC I is the most flexible receptor, while TL4 is the least flexible. This suggests that the vaccination complex may have a more dynamic interaction with MHC I, which could be essential for the process of antigen presentation.

**Figure 6:**
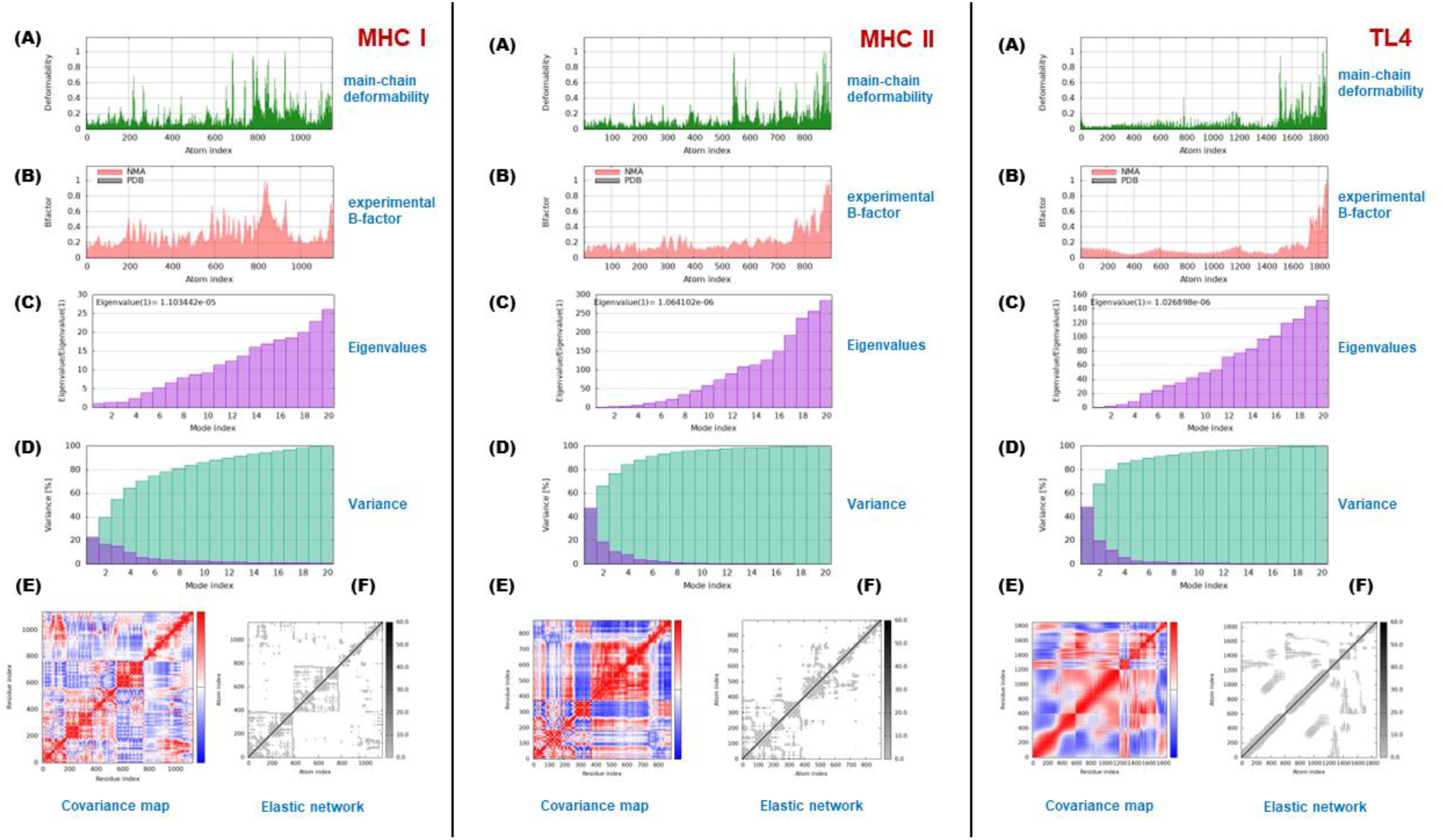
Molecular dynamics simulations (i-MODS) outputs of the vaccine complexes with three receptors MHC I, MHC II and TL4. **(A)** main chain deformability **(B)** experimental B-factor **(C)** Eigenvalues **(D)** variance **(E)** covariance map and **(F)** elastic network

### 3.7 Immune response simulation studies

The study on the vaccine’s immune simulation was carried out using the C-ImmSimm server, which predicts the activation of adaptive immunity and the immune interactions between epitopes and their specific targets [66]. The graph in **Figure 7(A)** categorizes antibodies based on isotype. Antibody levels (IgM and IgG) increase slowly following exposure to the antigen. This suggests that the immune system is starting to react to the antigen. Later on, the IgG antibody level becomes higher than the IgM antibody level. There is a shift in the immune response from mainly IgM to mostly IgG. This is a common feature of the immune response. The IgM antibody is typically the first immune response to an infection. Although the molecule demonstrates a robust capacity for binding to pathogens, it falls short in its ability to activate the complement system or opsonize pathogens for phagocytosis. IgG is the main antibody found in the blood. This molecule is smaller than IgM, but it is more effective in activating the complement system and enhancing the ability to engulf pathogens for phagocytosis. Moreover, IgG shows superior effectiveness in neutralizing toxins and viruses. The immune response shifts from an initial IgM-dominated phase to a subsequent IgG-dominated phase, as shown by the higher IgG antibody titer compared to the IgM antibody titer at later time points. This represents a common feature of the immune response. **Figure 7(B)** displays the B cell population per entity-state over time in days, with the quantity of B cells per mm³ on the vertical axis. The graph displays four states: active, internalized, duplicating, and anergic. Throughout the simulation, active B cells consistently maintain a population of around 400 cells/mm³, making them the most populated state. The B cells that are duplicating show a similar pattern to the active B cells, but at a lower level of around 300 cells/mm³. On day 15, the number of internalized B cells is notably lower than the number of active and duplicating B cells, reaching around 75 cells/mm³. Consistently, anergic B cells maintain a population of around 50 cells/mm³ throughout the simulation, making it the least populated state. **Figure 7(C)** shows the changes in B lymphocyte sub-populations over time, potentially due to a GBM vaccination. On the vertical axis, you can see the cell count per mm³, and on the horizontal axis, time in days is shown. The B cell count is increasing, as shown by the blue line. Suggesting that the GBM vaccine is enhancing B cell generation. Memory B cells experience a peak in numbers at first, then gradually decrease over time. These B cells have encountered a specific antigen in the past. The elevated initial number suggests that the vaccine is activating memory B cells. The decreasing trend suggests a transition of memory B cells into plasma cells or cell death. Various B cell types: This graph illustrates three B cell isotypes: IgM, IgG1, and IgG2. B cells have the ability to generate different types of antibodies. The concentration of all three isotypes has been observed to increase over time, reflecting the trend observed in the total B cell count. This graph clearly shows how the GBM vaccine is boosting B cell production and activation, including memory B cells and naive B cells. This suggests that the vaccine is promoting the production of all three main B cell types: IgM, IgG1, and IgG2. Displayed in **Figure 7(D)** is the number of CD4 T-helper lymphocytes per cubic millimeter on the y-axis, while the x-axis represents simulation time in days. This graph illustrates four different states of the CD4 T-helper lymphocytes: active, resting, anergic, and duplicating. This graph clearly shows a sharp rise in the quantity of active CD4 T-helper lymphocytes at first, which then levels off around 10 days later. CD4 T-helper lymphocytes experience an initial rapid increase, followed by a decline after around 10 days. CD4 T-helper lymphocytes become anergic gradually over time. The count of CD4 T-helper lymphocytes increases rapidly at first, then levels off by day 10. The graph clearly shows that the simulation anticipates a strong immune response to the vaccine design. There is a clear rise in active CD4 T-helper lymphocytes. Illustrated in **Figure 7(E)** are the CD4 T-helper lymphocytes count, which includes both total and memory cells following immune stimulation. The y-axis shows the cell count per mm³, while the x-axis indicates time in days. The graph shows two lines: one representing the total number of CD4 cells (TH not Mem) and the other showing the number of memory T cells (TH Mem). There is a consistent upward trend in both total and memory CD4 T cell counts over the course of the simulation, with the rate of increase slowing down over time. Deciphering the precise timing and values of peak cell counts from the graph can be quite difficult. It is clear that memory T cell count consistently exceeds the total cell count at all indicated time points. The graph demonstrates that the immune stimulation protocol successfully enhances CD4 T-helper lymphocytes, encompassing both total and memory cells. **Figure 7(F)** displays the CD4 T-regulatory lymphocytes count per state over time. On the graph, the vertical axis shows the cell count per cubic millimeter, and the horizontal axis indicates time in days. The graph illustrates a consistent rise in the overall count of CD4 T-regulatory lymphocytes (TR), encompassing both memory (TR Mem) and non-memory (TR not Mem) cells, as time progresses. In addition, there has been an increase in both active and resting T regulatory cells, with a greater number of resting cells. **Figure 7(G)** shows the number of CD8 T-cytotoxic lymphocytes per cubic millimeter (cells/mm³) on the y-axis. These immune cells possess the capacity to eradicate cancer cells. The x-axis shows the time that has passed since the vaccination. This graph highlights two main trends. With time, the number of active CD8 T-cytotoxic lymphocytes increases. It suggests that the vaccine is stimulating the immune system to boost the production of these cells. With time, there is a decrease in the number of anergic CD8 T-cytotoxic lymphocytes. Immune cells do not operate normally in a state of anergy. It suggests that the vaccine is helping to improve the function of these cells. Illustrated in **Figure 7(H)** is the concentration of CD8 T-cytotoxic lymphocytes (TC cells) over time, potentially in response to a vaccine or immune stimulation. Displayed on the vertical axis is the number of TC cells per cubic millimeter (mm³), with time in days shown on the horizontal axis. On the graph, you’ll see two lines labeled as "Total" and "TC Mem (y2)". Over time, the total number of TC cells has been observed to increase, with fluctuations in the number of memory T cells. As per **Figure 7(I)**, the concentrations of most cytokines and interleukins exhibit a rising pattern as time progresses. IL-10, IL-4, TGF-β, and IFN-β reach their peak levels around day 15 before stabilizing. Several cytokines peak around day 20, including IL-12, IL-2, TNF-α, IL-6, IL-18, and IL-23. The graph illustrates a steady rise in IFN-γ and IL-D levels throughout the 35-day period. Displaying the population per state on the y-axis and time in days on the x-axis is shown in **Figure 7(J)**. Here is a theoretical population trend of dendritic cells (DCs) over time. Every color represents a different state of the DCs: Total, Active, Resting, Internalized, Presenting-1, and Presenting-2. The graph shows a steady rise in the overall number of DCs as time progresses. Although the graph doesn’t show the state with the highest number of DCs, it effectively demonstrates the distribution of active, resting, and internalized DCs throughout the simulation. From days 0 to 10, there is a steady increase in DCs in the "Presenting-1" state. **Figure 7(K)** displays the count of Natural Killer (NK) cells per cubic millimeter (mm3) on the y-axis, with time in days on the x-axis. The figure illustrates a steady rise in the quantity of NK cells in the simulation. It is indicated that the GBM vaccine design being simulated effectively boosts the immune system to enhance NK cell production. Illustrated in **Figure 7(L)** is the total count of macrophages in four different states (resting, active, internalized, and presenting-MHC II) over a 35-day period. It is uncertain which state has the highest number of macrophages because the y-axis lacks a scale. It appears that the number of macrophages is increasing in all four states over time. Only a few macrophages (purple line) uptake and display MHC II in comparison to the overall macrophage population (blue line). Resting macrophages consistently have higher numbers than active macrophages.

**Figure 7:**
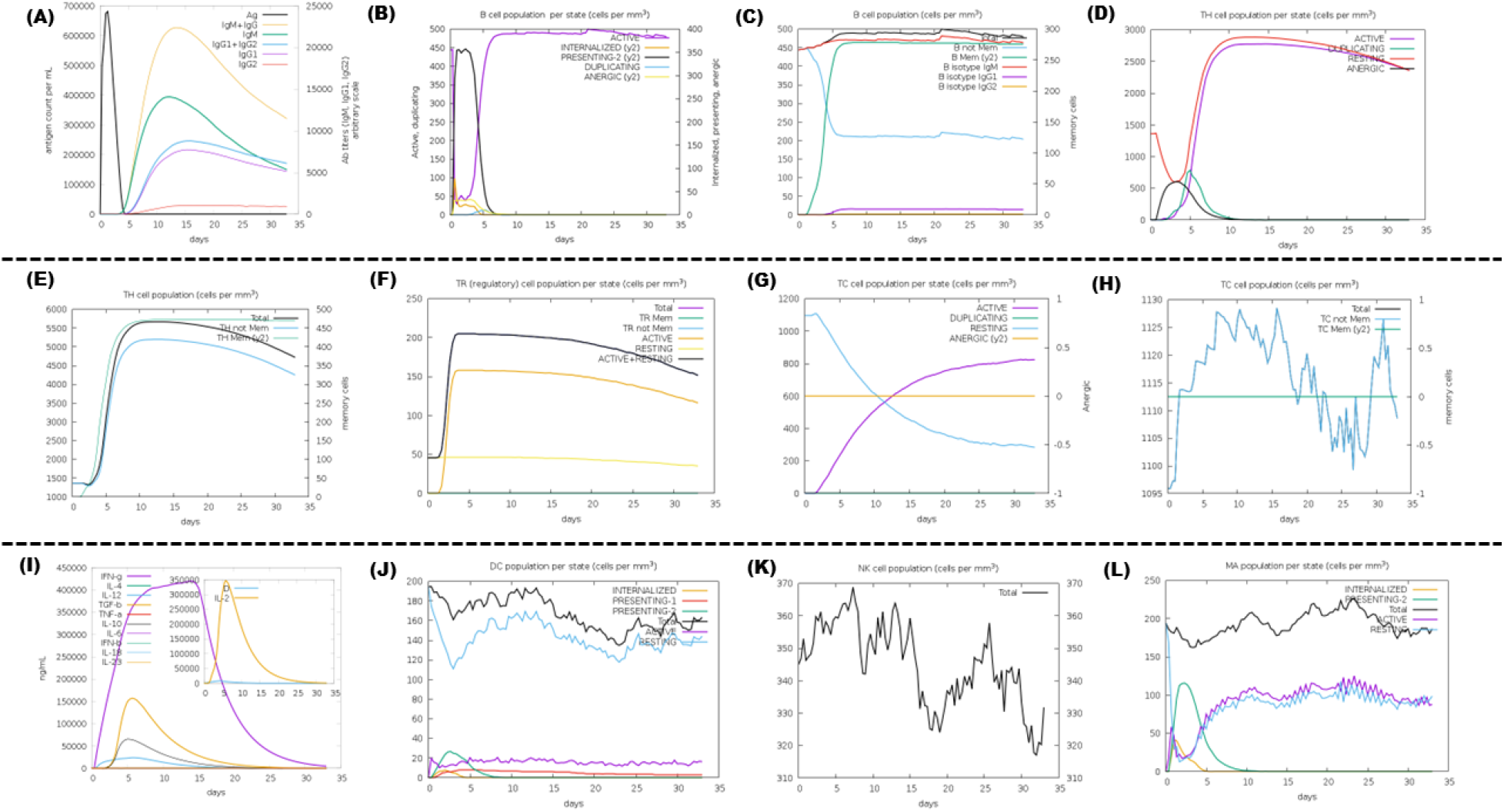
Immune response simulation studies **(A)** Antigen and immunoglobulins. Antibodies are classified based on their isotype. **(B)** The population of B lymphocytes per entity-state is displayed, indicating the counts for active presentation on class-II internalized antigens, duplication, and anergy. **(C)** The total count of B lymphocytes, as well as their memory cells, are further divided into different isotypes such as IgM, IgG1, and IgG2. **(D)** The count of CD4 T-helper lymphocytes is divided into different categories based on their state, including active, resting, anergic, and duplicating. **(E)** Count of CD4 T-helper lymphocytes. The plot displays the total and memory counts. **(F)** The quantification of CD4 T-regulatory cells. This graphic displays the total, memory, and per entity-state counts. **(G)** The quantification of CD8 T-cytotoxic lymphocytes per entity-state. **(H)** Count of CD8 T-cytotoxic lymphocytes. Displayed are the total and memory information. **(I)** Concentration of cytokines and interleukins. The danger signal is indicated by D in the inset plot. **(J)** Dendritic cells Antigenic peptides can be presented by DC on both MHC class-I and class-II molecules. **(K)** Total count of Natural Killer cells **(L)** The total count of internalized molecules is observed on both active and resting macrophages expressing MHC class-II.

### 3.8 Reverse transcription, Codon adaptation and in-silico cloning

The EMBOSS Backtranseq tool was used to perform reverse transcription of the vaccine peptide sequence into a nucleic acid sequence. Subsequently, the JCat server was employed to optimize it. The CAI value of the pasted sequence is 0.3057553005583999 and the GC content of the pasted sequence is 69.009009009009. Post-optimization, the CAI-Value of the enhanced sequence is 1.0, while the GC-Content is 54.234234234234236, both of which lie within the required range. The nucleic acid sequence was optimized and consisted of 1110 nucleotides. The results of codon adaptation are illustrated in **Table 5**. **Figure 8** illustrates the plasmid map resulting from the in-silico cloning of a GBM vaccine DNA fragment into a pET-28a(+) vector using SnapGene software. The initial pET-28a(+) plasmid measures 5028 base pairs (bp). The DNA fragment inserted into the GBM vaccine is 1110 base pairs long. BlpI and BsshII restriction enzymes were employed to cleave the plasmid and insert the fragment. The enzyme cleavage sites are located at 80 bp for BlpI and 1534 bp for BsshII. The lacI gene and lac promoter are located on the plasmid backbone. These components play a role in controlling the expression of the inserted gene. The Kanamycin resistance gene (KanR) is located on the plasmid backbone. This enables the selection of clones that have effectively integrated the plasmid. The region where the GBM vaccine fragment was inserted is known as the MCS, located between the BlpI and BsshII restriction sites. The fragment is bordered by a 6xHis tag, which can aid in purifying the protein.

**Figure 8:**
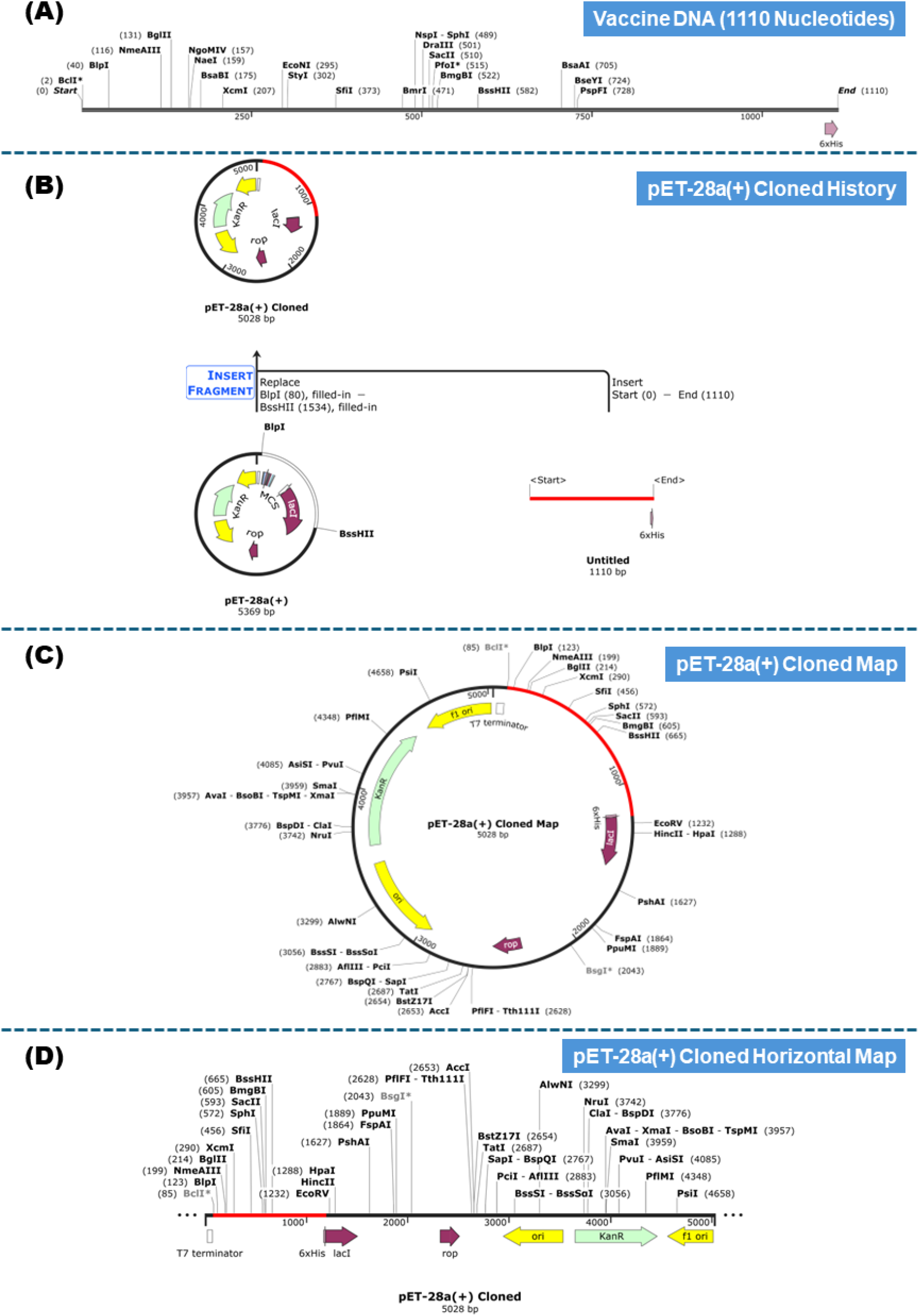
In silico cloning of the vaccine using pET-28(+) vector **(A)** The nucleic acid sequence of the vaccine **(B)** Cloning history **(C)** Cloned map **(D)** Horizontal map of the cloned vector.

**Table 5:**
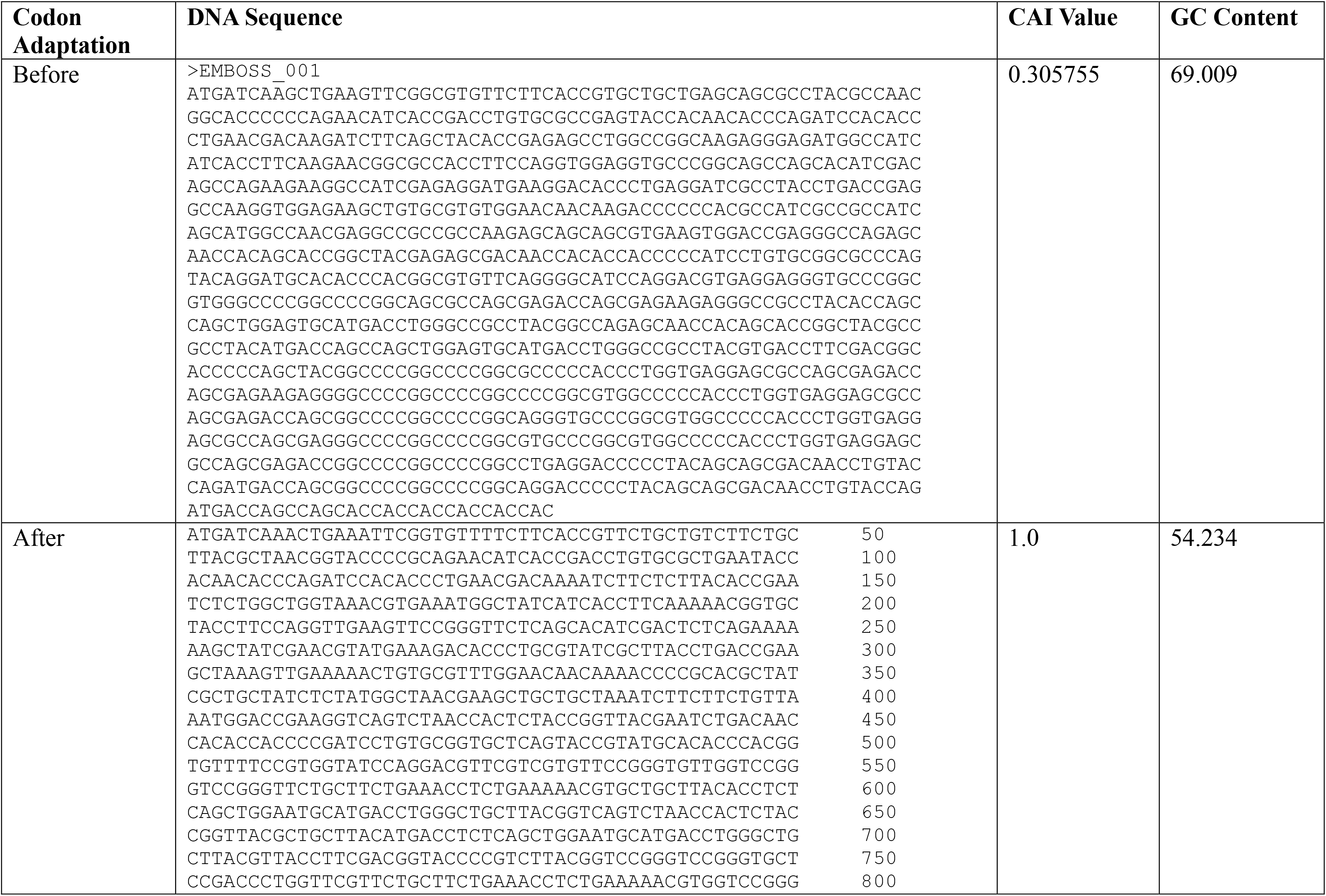
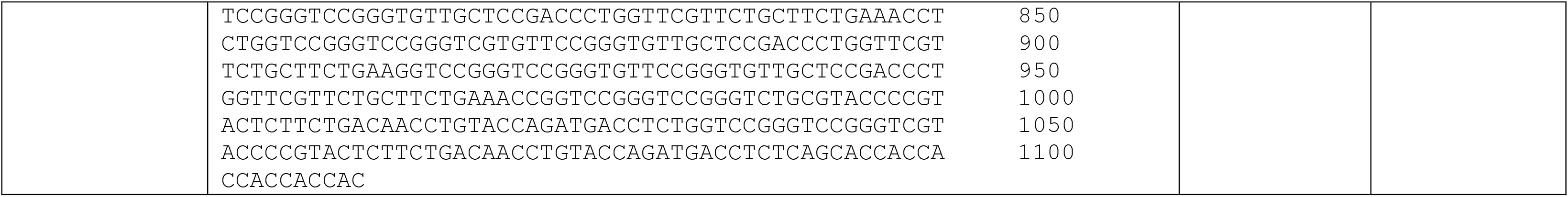
Nucleic acid sequence and codon adaptation.

## 4. Discussion

The aim of this study was to investigate the potential use of computational approaches to construct a multi-epitope vaccine targeting the WT1 protein for GBM immunotherapy. By identifying immunogenic epitopes from WT1 and evaluating their predicted antigenicity and allergenicity, immuno-informatics approaches were to be used to design a multi-epitope vaccine which are predicted to be non-toxic and non-allergic. The process comprised a thorough analysis that began with the protein sequences for WT1 being retrieved. Later, B-cell and T-cell epitopes were predicted using well-established prediction techniques. The antigenicity and allergenicity of the predicted epitopes were assessed. Ensuring that the immune response targets the tumor exclusively while minimizing any potential adverse effects was the main goal of this evaluation. Additional predictions assessed the epitopes’ capacity to induce cytokines, which are essential for immune system activation. To optimize their display on antigen-presenting cells (APCs), which in turn promotes T-cell identification, the MHC-binding epitopes were grouped with B cell epitopes. After the selected epitopes were combined with the proper adjuvants and linkers, a vaccine construct was produced. The construct’s physicochemical characteristics were carefully examined to make sure it was stable, soluble, and feasible to express. Accurate predictions of the vaccine construct’s secondary and tertiary structures were made using computational modelling methods. The structural predictions shed important light on the vaccine’s general shape and its interactions with immune receptors. The tertiary structure of the model was refined and validated to increase its accuracy even further. Molecular docking simulations were used to evaluate the vaccine construct’s binding affinity to the MHC I, MHC II and TLR4 receptors, an essential part of the innate immune system that triggers an immunological response. Important insights into the long-term stability and dynamics of the vaccine-receptor combination were obtained from the molecular dynamics simulations. To enhance its expression in a selected host system, the vaccine sequence underwent a process of codon adaptation and back-translation, which will facilitate future research in lab and living creatures. Through in-silico cloning simulations, the vaccine construct’s integration into an appropriate expression vector was further validated, indicating its practicality. To predict how the vaccine under consideration might activate different immune cells, a simulation was run. The significance of applying immuno-informatics techniques to expedite vaccine development is underscored by this study. The benefits of designing vaccines with multiple epitopes are numerous. By focusing on distinct WT1 epitopes, it may be possible to strengthen the immune system and address tumor heterogeneity, a frequent characteristic of GBM. Adjuvants can significantly increase the vaccine’s immunogenicity. The strategic selection of adjuvants and epitopes is made possible by immuno-informatics tools, which facilitate a focused and efficient approach which lays the groundwork for future experiments; preclinical: in-vitro studies, in-vivo and clinical studies to validate the vaccine design reported in this paper. Although immuno-informatics techniques cannot perfectly duplicate the intricate workings of the human immune system, they do offer valuable insights. To confirm the immunogenicity and effectiveness of the developed vaccine construct, in-vitro and in-vivo studies must be carried out. To determine the best dosage, mode of administration, and any adverse effects of the vaccine, pre-clinical and clinical trials are crucial. An additional major problem is the immunosuppressive microenvironment of GBM tumors. Combining immunosuppressive methods with the WT1-targeted vaccine may be important to obtain a strong and long-lasting anti-tumor response. The emphasis on just one tumor antigen, WT1, is another drawback. Although targeting WT1 has obvious potential, other tumor-associated antigens may need to be included in the vaccine design to elicit a more thorough immune response.

## 5. Conclusion

An immuno-informatics-driven multi-epitope vaccine for glioblastoma multiforme (GBM) that specifically targets WT1 represents a promising groundwork for future research in addressing the challenges presented by this highly malignant brain tumor. GBM continues to pose a significant challenge in the field of oncology, as there are limited treatment options available and patient outcomes are often discouraging. The development of innovative immunotherapeutic strategies, such as the one proposed in this in-silico study, might facilitate the search for the novel treatment of GBM. WT1 is being focused upon because it is a promising tumor-associated antigen and is expressed at higher levels in GBM tissues than in normal cells. This work identified potential B and T-cell epitopes by doing a comprehensive analysis of the WT1 protein using immuno-informatics methods. This study serves as a basis for the development of a multi-epitope vaccination. The vaccine design, which combines B-cell and MHC-binding epitopes with adjuvants for enhanced immunogenicity, has been predicted to elicit a strong and targeted anti-tumor immune response. When compared to conventional methods, the multi-epitope vaccine design offers several advantages. First, by focusing on several epitopes, it tackles the difficulties associated with tumor heterogeneity and may reduce the likelihood of immune escape mechanisms. Moreover, adjuvants improve antigen presentation and activate immune cells, fostering an environment that facilitates the formation of a potent anti-tumor response. Furthermore, the use of immuno-informatics speeds up the development of vaccines by facilitating rapid epitope screening and thoughtful vaccine construct design. Comprehensive investigation was carried out by the study, which included simulations of immune responses, structural elucidation, physicochemical property simulation assessment, and molecular docking. These results provide a solid basis for upcoming preclinical and clinical research. Confirming the safety, efficacy, and possible use of the suggested vaccine candidate will require completing in vitro can be used to evaluate the efficacy of the vaccine by looking at how well it stimulates immune cells and causes cytotoxicity against GBM cells. Furthermore, in vivo studies carried out on animal models could provide important insights into the immunogenicity and therapeutic efficiency of the vaccine. Furthermore, further investigation could be directed in several different ways towards optimizing and refining the multi-epitope vaccine that specifically targets WT1. This might entail optimizing vaccine delivery strategies to increase immune response and guarantee accurate targeting of tumor sites, enhancing vaccine composition by selecting the best epitopes and adjuvants, and researching combined strategies with other immunotherapies like immune checkpoint inhibitors or adoptive cell therapies. To maximize therapeutic outcomes, it is also critical to concentrate on finding predictive biomarkers of vaccine response and creating plans to combat immunosuppressive mechanisms in the tumor microenvironment. The potential to improve patient outcomes and revolutionize GBM treatment approaches underscores the significance of continued research in this sector, even though the journey from bench to bedside can be difficult. For GBM sufferers who are battling a fatal disease, there may be new hope thanks to the novel vaccination idea. There is a lot of potential for its application in clinical practice.

## Declaration Statements

### Competing Interests

The authors have no relevant financial or non-financial interests to disclose.

### Author Contributions (CRediT)

Conceptualization: **T.A.**, Methodology: **T.A.**, Software: **T.A.**, Formal analysis: **T.A.**, Investigation: **T.A.**, Data Curation: **T.A.**, Writing - Original Draft: **T.A.**, Writing - Review & Editing: **T.A.**, Visualization: **T.A.**

## Abbreviations

GBM: Glioblastoma Multiforme
WT1: Wilms’ Tumor 1
MHC: Major histocompatibility complex
CTL: Cytotoxic T lymphocytes
HTL: Helper T lymphocyte
HLA: Human leukocyte antigens
DC: Dendritic cells
FASTA: Fast Adaptive Shrinkage Threshold Algorithm
IFN-γ: Interferon-gamma
GRAVY: Grand average of hydropathicity index
TLR4: Toll-like receptor 4
IgM: Immunoglobulin M
IgG: Immunoglobulin G
CD4: Clusters of differentiation 4
TR: T-regulatory lymphocytes
CD8: Cluster of differentiation 8
TGF-β: Transforming growth factor-beta
IFN-β: Interferon beta
TNF-α: Tumor Necrosis Factor alpha
NK: Natural Killer
MCS: Multiple cloning site
APC: Antigen-presenting cell

## Supplementary Information

Supporting Information file is available via DOI 10.5281/zenodo.10890996 or (https://zenodo.org/records/10890997)

---

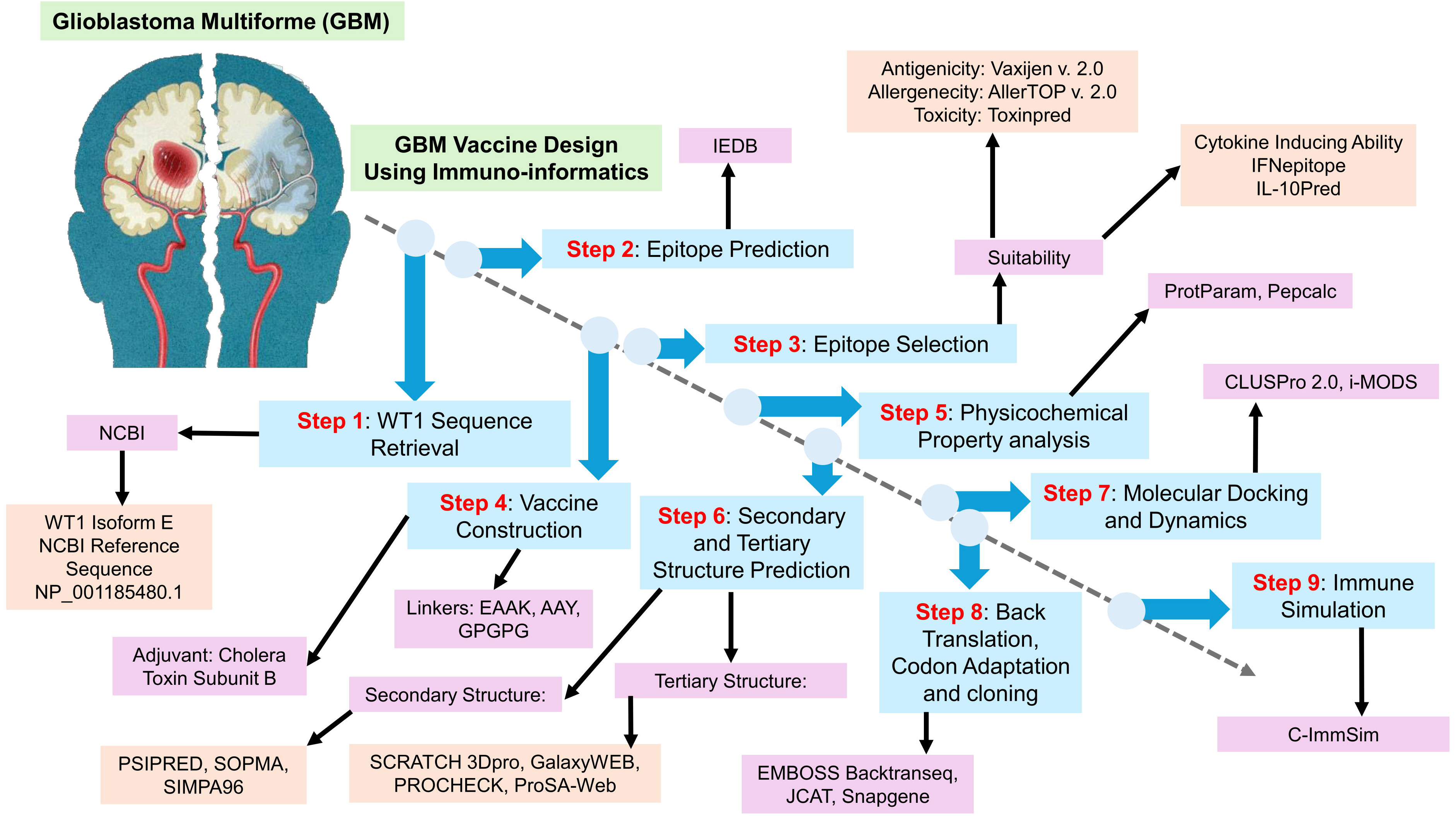

## Notes

https://zenodo.org/records/10890997

### Competing Interest Statement

The authors have declared no competing interest.

